# Differential Alterations of Gut Microbiota in Girls with Isolated Obesity, Isolated Precocious Puberty, and Their Comorbidity

**DOI:** 10.64898/2025.12.10.693599

**Authors:** Han Xuan, Hou Yangnan, Su Ganglin, Zhang Zhaoxia, Zhang Qin, Dong Guoqing

## Abstract

**Background:** Childhood obesity and precocious puberty (PP) are escalating public health concerns with a significant comorbidity. Emerging evidence implicates gut microbiota dysbiosis in both conditions, yet microbial associations with their co-occurrence remain unexplored.

**Methods:** A total of 68 girls (18 normal, 25 PP-only, 18 obesity-only, and 7 obesity-PP comorbidity) were enrolled. 16S rRNA sequencing was applied to determine gut microbiota of girls. Microbial diversity, composition, and differential abundant taxa were analyzed.

**Results:** While overall microbial diversity and community structure were similar across groups, obesity-only subjects exhibited significantly increased microbial richness (Chao1 index, P<0.01 vs. all groups), characterized by a higher Firmicutes/Bacteroidota ratio (P<0.001 vs. normal/PP), increased Proteobacteria (P<0.001 vs. normal), and enrichment of genera including Blautia, Faecalibacterium, Roseburia, Anaerostipes, Haemophilus, and Lachnospiraceae_NK4A136_group (P<0.05 vs. other groups). PP-only subjects showed reduced Verrucomicrobiota (P<0.001 vs. normal/obesity) and elevated Escherichia_Shigella (P<0.05 vs. normal). Most strikingly, the obesity-PP comorbid group exhibited a unique gut microbiota profile characterized by a dramatic depletion of Firmicutes compared to the obesity-only group (P<0.001) and a remarkable nearly 100-fold increase in Verrucomicrobiota abundance (P<0.001 vs. other groups), suggesting a distinct microbial ecotype potentially contributing to the pathophysiology of this comorbidity.

**Conclusion:** This study delineated unique gut microbiota alterations associated with isolated obesity, isolated PP, and their comorbidity in girls. The distinct dysbiotic pattern in comorbid obesity-PP suggested complex microbiota-host interactions, providing a new microbiological perspective for understanding the mechanism of comorbidity between obesity and precocious puberty.

## 1. Introduction

The escalating prevalence of both childhood obesity and precocious puberty (PP) has emerged as parallel public health crises. Data from the Chinese National Surveys on Students’ Constitution and Health reveal a dramatic rise in obesity prevalence among children and adolescents aged 7–18 years, increasing from 0.1% in 1985 to 9.6% in 2019—a 75.6-fold surge (Yuan et al., 2024). PP, a disorder characterized by premature development of secondary sexual characteristics, has shown concerning trends in China. A 2009–2010 survey of 18,707 elementary and middle-school students across six cities reported an overall PP prevalence of 0.43%, with higher rates in girls (0.48%) than boys (0.38%) (Zhu et al., 2013). More recently, Liu et al. found that the adjusted PP prevalence among grades 1–3 students in Zhongshan, Guangdong reached 6.19% (girls: 11.47%; boys: 3.26%) when assessed by Tanner staging (Y. F. Liu et al., 2021). Both conditions are associated with serious health consequences, including metabolic syndrome, type 2 diabetes (Shi et al., 2022), and psychological comorbidities such as anxiety, depression, and behavioral problems (Huang et al., 2021; Iwatate et al., 2023; Nacinovich et al., 2016; Small & Aplasca, 2016; Xie et al., 2023).

Notably, childhood obesity and PP frequently co-occur. Studies indicate that girls with obesity have a 1.78-fold higher risk of PP compared to their normal-weight peers (G. Y. Liu et al., 2021). Genetic evidence further supports this connection, as genome-wide association studies (GWAS) have identified BMI-increasing alleles that also correlate with earlier menarche timing (Perry et al., 2014), further confirming the comorbidity of childhood obesity and PP from the perspective of genetics. These findings underscore the importance of identifying shared etiological factors between obesity and PP to develop effective interventions for pediatric populations.

Emerging research suggests that the gut microbiota, a complex ecosystem of trillions of microorganisms in the gastrointestinal tract, plays a crucial role in metabolic regulation, hormone balance, and immune function, potentially influencing both conditions. Specifically, gut microbes influence obesity progression by regulating energy harvest through enhanced production of short-chain fatty acids, modulating fat storage via bile acid metabolism and lipoprotein lipase activity, and promoting low-grade inflammation through mechanisms like lipopolysaccharide (LPS)-induced insulin resistance (Angelini et al., 2024; Cani & Van Hul, 2024; Gomes et al., 2018; L. Zhang et al., 2024). Furthermore, certain gut microbiota influence the release of intestinal hormones such as GLP-1, ghrelin, PYY, and leptin, thereby modifying signaling along the gut-brain axis that regulates satiety and feeding behavior via hypothalamic pathways (Torres-Fuentes et al., 2015). Meanwhile, emerging evidence suggests that gut microbiota can directly affect precocious puberty or indirectly through obesity. Animal studies using fecal microbiota transplantation from donors with precocious puberty (either human patients or rat models) to female recipient rats showed earlier puberty onset, which was associated with increased serum LH, FSH, and estradiol levels, along with heightened hypothalamic expression of Kiss1 and Gnrh genes (Bo et al., 2022; Qian et al., 2024). Remarkably, microbial reconstitution can reverse maternal high-fat diet-induced precocious puberty in mice (Wang et al., 2020), while microbiota-derived mediators (leptin, kisspeptin, and neuroexcitation peptides) may underlie the obesity-PP comorbidity (Kang et al., 2018). These findings collectively indicate that targeted interventions modulating gut microbiota composition and function may represent a novel therapeutic approach for preventing and managing childhood obesity and its associated endocrine disorders, particularly PP.

Collectively, these findings suggest that gut microbiota may influence obesity and precocious puberty through both direct mechanisms and indirect pathways mediated by obesity. Although accumulating evidence has demonstrated alterations in gut microbial composition in childhood obesity (Fei & Zhao, 2013; Hjorth et al., 2019; Mathur & Barlow, 2015; L. Wang et al., 2024) and precocious puberty (Bao et al., 2025; Calcaterra et al., 2022; L. Wang et al., 2024; Yue & Zhang, 2024) separately, studies investigating the microbial associations with both conditions simultaneously remain limited. To address this critical knowledge gap, we conducted the present study. Employing a strategic grouping design to control for potential sex confounding effects, we performed a cross-sectional comparison of gut microbiota profiles among four distinct groups: obesity alone, precocious puberty alone, comorbid obesity with precocious puberty, and normal control girls. This study aims to investigate whether gut microbiota is associated with the co-occurrence of obesity and precocious puberty, with the ultimate goals of identifying unique microbial signatures that may characterize this comorbid condition and providing scientific evidence for developing preventive and therapeutic strategies against these childhood disorders.

## 2. Materials and methods

### 2.1 Study population

The present study enrolled study subjects at the Longhua Branch of Shenzhen People’s Hospital, Shenzhen from January, 2022 to August 2023. The children with precocious puberty were identified based on clinical diagnostic examination and diagnosed by professional pediatricians. The obesity was diagnosed by body mass index (BMI) according to the diagnostic criteria of Working Group on Obesity in China (Li et al., 2010). After obtaining the informed consent of the parents, we collected stool samples of the children to determine their status of gut microbiota. The exclusion criteria were as follows: (1) a diagnosis of intestinal disorders (e.g., inflammatory bowel disease) or use of medications influencing gastrointestinal motility (e.g., stool softeners); (2) current participation in structured dietary therapy, dietary interventions, or professionally supervised complementary and alternative therapies; (3) recent (within one month) use of probiotics, antibiotics, antifungals, anti-inflammatory agents, or antioxidants. Finally, a total of 68 girls (average age: 7.20 ± 1.03) in the hospital outpatient department was recruited as research subjects for this study, including 18 normal girls, 25 girls with only precocious puberty, 18 girls with only obesity, and 7 girls who were obesity-PP comorbidity (both obese and precocious puberty).

#### Ethical approval and informed consent

This study was reviewed and approved by the Scientific Research Ethics Committee of Shenzhen People’s Hospital (Approval No. LL-KY-2023096-02). The study was conducted in accordance with the ethical principles of the Declaration of Helsinki and relevant national regulations. Prior to participation, written informed consent was obtained from the legal guardians of all children involved in this study.

### 2.2 Sample collection and extraction of DNA

Stool specimens were collected using sterile fecal containers, immediately placed on ice, and stored at −80°C within 30 minutes. Microbial DNA was extracted using the E.Z.N.A.® Soil DNA Kit (Omega Bio-tek, Norcross, GA, USA) following the manufacturer’s instructions. DNA concentration and purity were assessed using a NanoDrop 2000 UV-vis spectrophotometer (Thermo Scientific, Wilmington, USA), and integrity was verified by 1% agarose gel electrophoresis.

### 2.3 16S rRNA gene amplicon sequencing

The V3-V4 hypervariable regions of the bacterial 16S rRNA gene were amplified via PCR using primers 338F (5′-ACTCCTACGGGAGGCAGCAG-3′) and 806R (5′-GGACTACHVGGGTWTCTAAT-3′) on a GeneAmp 9700 thermocycler (ABI, USA). The PCR conditions consisted of initial denaturation at 95°C for 3 min; 27 cycles of 95°C for 30 s, 55°C for 30 s, and 72°C for 45 s; and a final extension at 72°C for 10 min. Reactions were performed in triplicate using a 20 μL mixture containing 4 μL of 5× FastPfu Buffer, 2 μL of 2.5 mM dNTPs, 0.8 μL of each primer (5 μM), 0.4 μL of FastPfu Polymerase, 0.2 μL of BSA, and 10 ng of template DNA. PCR products were purified from a 2% agarose gel using the AxyPrep DNA Gel Extraction Kit (Axygen Biosciences, USA) and quantified with QuantiFluor™-ST (Promega, USA).

Purified amplicons were pooled in equimolar ratios and subjected to paired-end sequencing (2 × 300 bp) on an Illumina MiSeq platform (Illumina, San Diego, USA) by Majorbio Bio-Pharm Technology Co. Ltd. (Shanghai, China), following standard protocols.

### 2.4 Bioinformatics analysis, statistical analysis, and data visualization

The raw sequences were assigned to different samples based on their corresponding barcodes. Fastp software (v 0.20.0) was employed for adapter removal and low-quality sequence filtration. The filtered high-quality reads were then merged, and subsequently subjected to the DADA2 pipeline for removal of chimeric sequences. Operational taxonomic units (OTUs) table were then generated at a sequence similarity cutoff value of 97% using QIIME2. The taxonomical classification of the OTUs was assigned using the QIIME2 pre-trained naïve Bayes classifier, which was trained on the SILVA database (Release 138). Then, MicrobiomeAnalyst was employed for downstream analysis, encompassing the computation of good’s coverage and facilitating visualization as well as statistical examination of microbial communities at diverse taxonomic levels to discern dissimilarities in microbiomes and diversity across distinct groups.

The alpha diversity of microbial communities, as measured by the Chao1 and Shannon indices, and differences between groups were analyzed with the Kruskal-Wallis test. The beta diversity was estimated by calculating Bray-Curtis distance and visualized through the principal coordinate analysis (PCoA) method. The significance of the PCoA plot was assessed using permutational multivariate analysis of variance (PERMANOVA), which employs distance metrics to validate the strength and statistical significance of sample groupings. The differential taxonomy between the two groups was compared using statistical method EdgeR, with a false discovery rate (FDR) correction applied to control for multiple comparisons. Statistical significance was defined as an FDR-adjusted P value < 0.05.

## 3. Results

### 3.1 Composition of microbial community

In total, 68 girls (including 18 normal girls, 25 girls with precocious puberty, 18 obese girls, and 7 girls with both obesity and precocious puberty) were collected and analyzed in this study. After the quality optimization of the original data, the average number of valid sequences for normal, precocious, obesity-PP comorbidobesity-PP comorbid, and obese samples were found to be 6,234, 6,569, 6.872, and 25,972, respectively (Table. S1). In addition, the good’s coverage was over 99.9% for all samples (Table. S1), indicating that the samples were almost completely sampled, and sufficient sequencing to accurately estimate the gut microbiota in 68 fecal samples.

A total of 11 phyla were identified in 68 samples (Fig. 1A), with *Firmicutes* (average 51.66% ± 17.40%), *Bacteroidota* (38.73% ± 19.55%), *Actinobacteriota* (4.50% ± 6.66%), and *Proteobacteria* (3.83% ± 3.88%) were the predominant phyla, collectively accounting for 92.73%-100% of each sample, except for A339116448 (58.49%). At the genus level, the predominant genus included *Bacteroides* (27.69% ± 17.09%), *Prevotella* (9.31% ± 20.96%), *Faecalibacterium* (7.00% ± 5.72%), *Roseburia* (6.16% ± 6.68%), *Subdoligranulum* (4.66% ± 7.85%), *Bifidobacterium* (4.00% ± 6.37%), and *Blautia* (3.62% ± 2.32%) in all samples (Fig. 1B), and these seven genera collectively accounted for 28.97%-89.95% of each sample. Moreover, the bacterial composition of gut microbiota in different group was highly similar at the phylum (Fig. S1), however, the result showed that the obese group had a significantly different gut microbiome composition than the other three groups at the genus level (Fig. 2).

**Fig 1.**
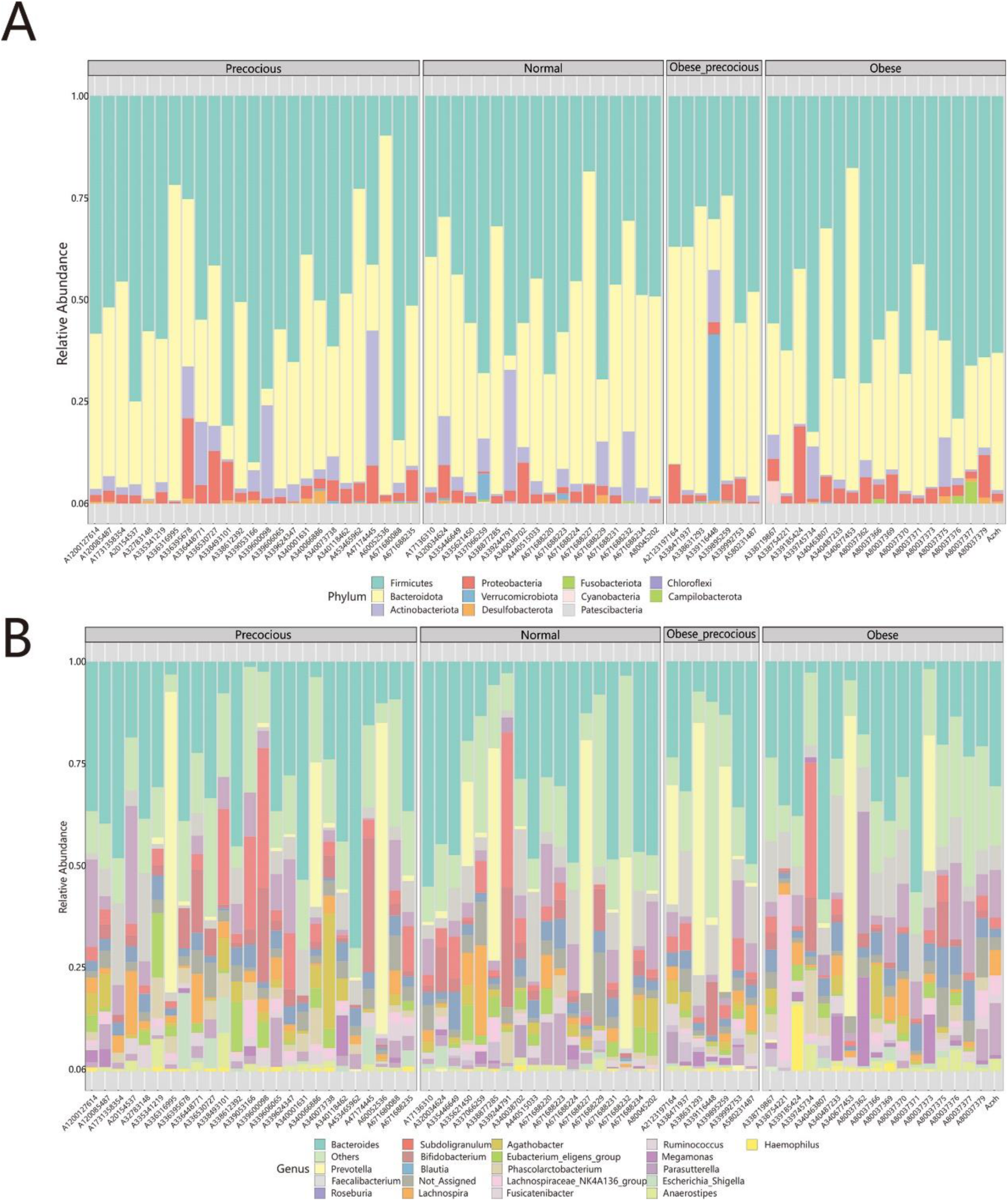
Gut microbiota comparison among various groups at (A) phylum level and (B) genus level.

**Figure 2.**
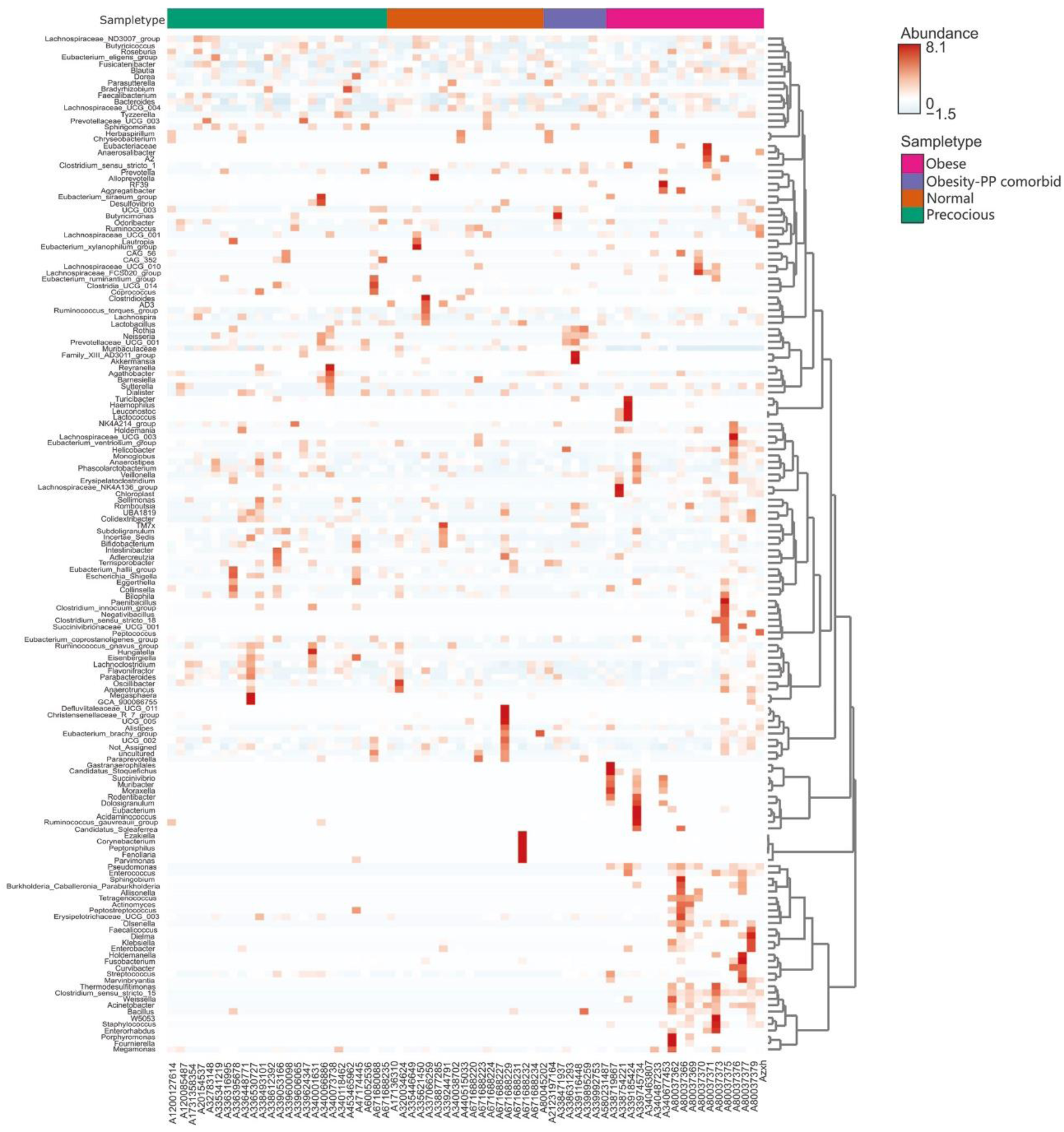
Gut microbiome composition of girls at the genus level.

### 3.2 Microbial community diversity in different groups

The microbial richness and diversity were represented by Chao1 and Shannon indices, respectively. Notably, the obese group exhibited significantly higher microbial richness compared to the normal group, the precocious group, and the obesity-PP comorbidobesity-PP comorbid group (Kruskal-Wallis, all *P* < 0.01), and there was no significant difference between the other groups, suggesting that obesity was significantly associated with increasement in the number of bacterial species, while precocious and obesity-PP comorbid did not change it significantly (Fig. 3A). However, there was no significant difference in microbial diversity between the four groups (Fig. 3B), indicating neither obese nor precocious causes significant changes in the diversity of gut microbiota. In addition, the β-diversity, as determined by PCoA analysis, revealed no significant differences in microbial structure were observed four groups (Bray-Curtis, PERMANOVA, *P* > 0.05) (Fig. 4).

**Figure 3.**
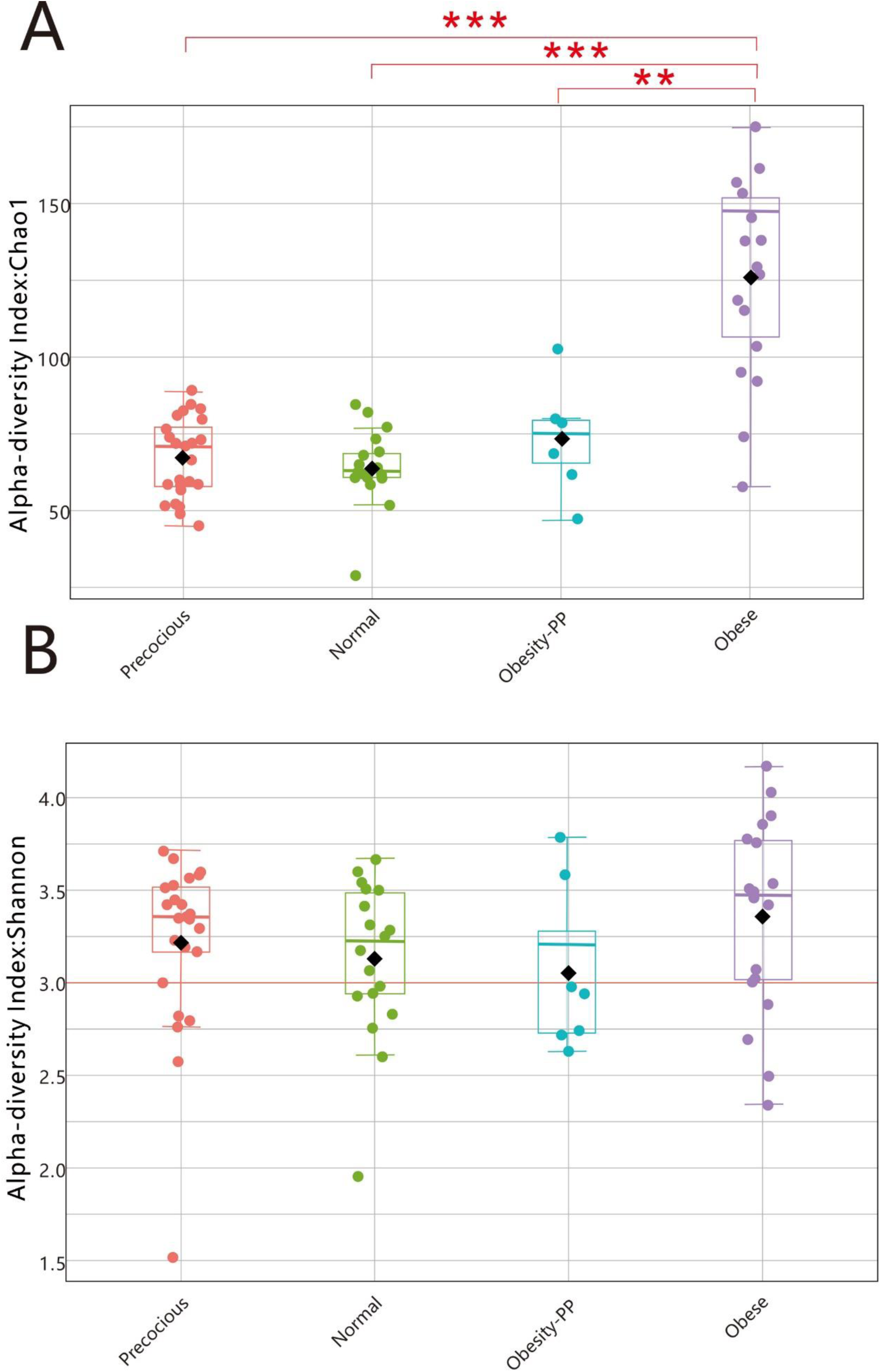
Comparison between microbial Alpha-diversity among different groups.

**Figure 4.**
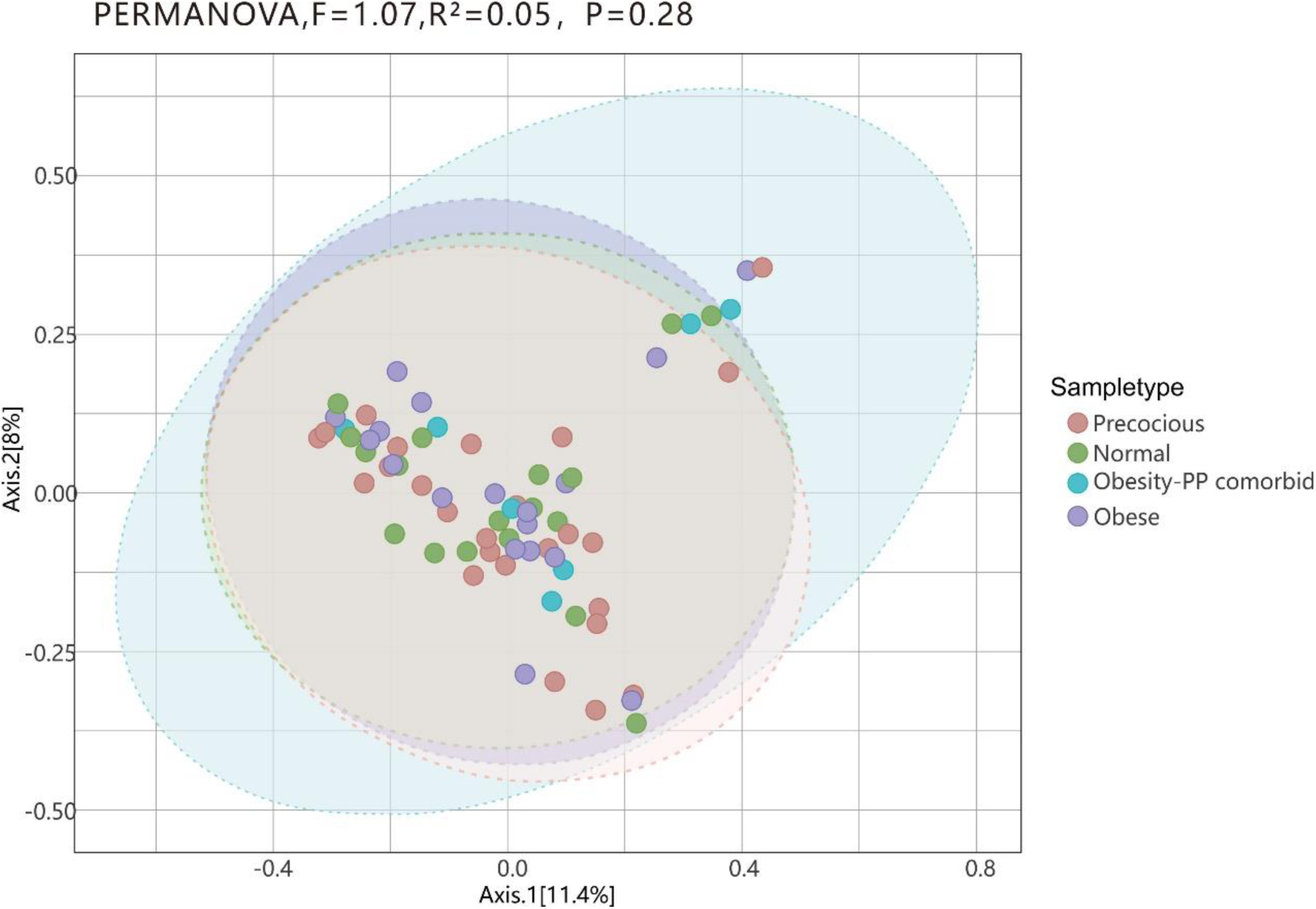
Comparison between microbial β-diversity among different groups.

### 3.3 Species with significant differences in different groups

Based on the relative abundance distribution of microbial communities at the phylum and genus levels, we employed statistical method EdgeR to identify species exhibiting statistically significant differences among different groups (Table 1). Specifically, there were 3, 2, 1, 8, 5, and 6 phyla in the in normal and precocious groups, normal and obesity-PP comorbid groups, precocious and obesity-PP comorbid groups, normal and obese groups, obesity-PP comorbid and obese groups, precocious and obese groups, respectively. In species exhibiting higher relative abundance (relative abundance over 1%), the abundance of *Firmicutes* and *Proteobacteria* in the obese group significantly higher than that in both normal group and precocious group (all *P* < 0.001); however, the abundance of Bacteroidota in the normal group and precocious group were significantly higher than that in the obese group (all *P* < 0.001). In addition, the abundance of *Firmicutes* in the obese group significantly higher than that in the obesity-PP comorbid group (*P* < 0.001). The abundance of *Verrucomicrobiota* in the obesity-PP comorbid group was significantly higher than that other three groups (all *P* < 0.001), and the normal group was significantly higher than that in the both precocious group and obese group (all *P* < 0.001).

**Table 1.**
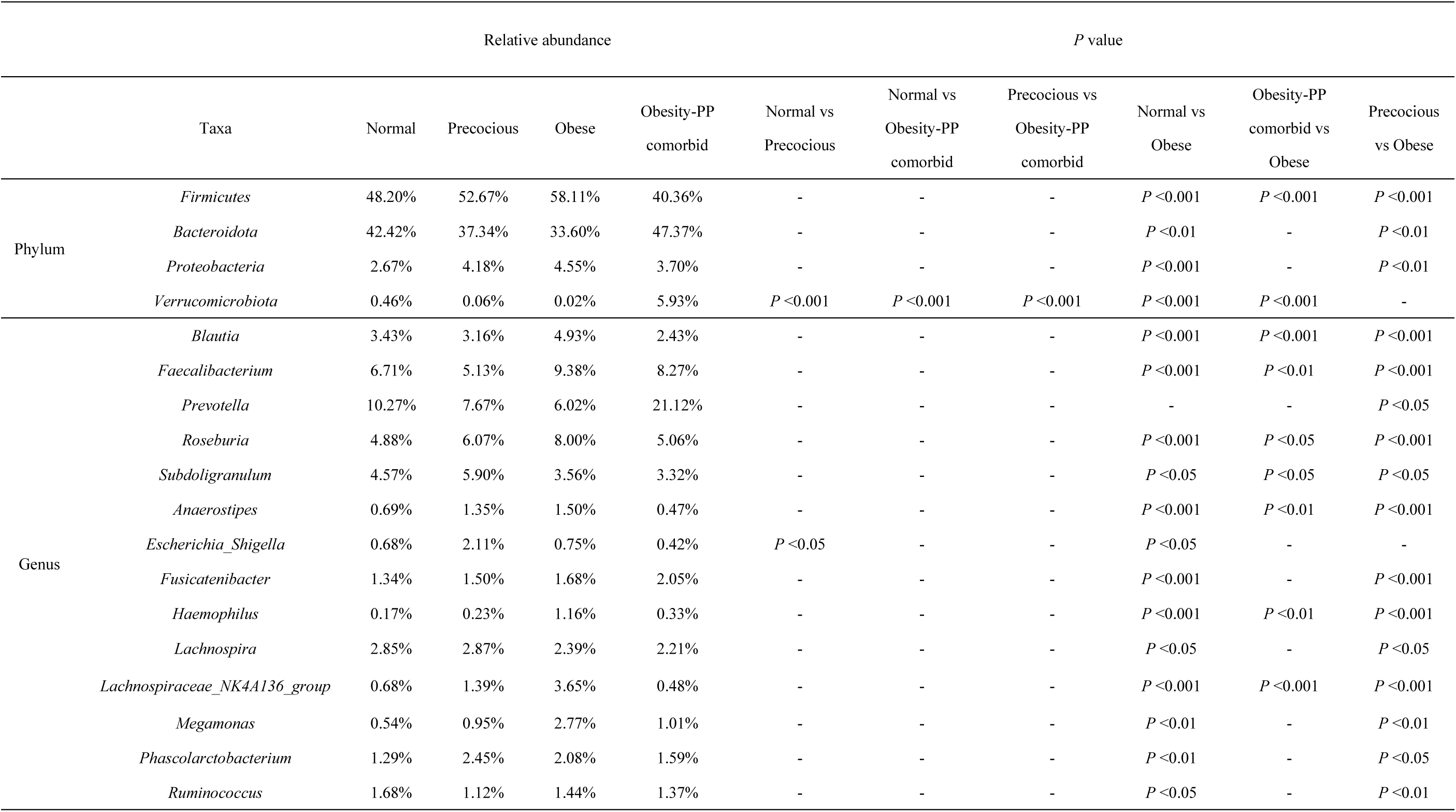
Taxa with significantly differential relative abundance and their relative abundance in the gut microbiome of four groups.

|  |  | Relative abundance |  |  |  | P value |  |  |  |  |  |
| --- | --- | --- | --- | --- | --- | --- | --- | --- | --- | --- | --- |
|  | Taxa | Normal | Precocious | Obese | Obesity-PP<br>comorbid | Normal vs<br>Precocious | Normal vs<br>Obesity-PP<br>comorbid | Precocious vs<br>Obesity-PP<br>comorbid | Normal vs<br>Obese | Obesity-PP<br>comorbid vs<br>Obese | Precocious<br>vs Obese |
| Phylum | *Firmicutes* | 48.20% | 52.67% | 58.11% | 40.36% | - | - | - | $P < 0.001$ | $P < 0.001$ | $P < 0.001$ |
| | *Bacteroidota* | 42.42% | 37.34% | 33.60% | 47.37% | - | - | - | $P < 0.01$ | - | $P < 0.01$ |
| | *Proteobacteria* | 2.67% | 4.18% | 4.55% | 3.70% | - | - | - | $P < 0.001$ | - | $P < 0.01$ |
| | *Verrucomicrobiota* | 0.46% | 0.06% | 0.02% | 5.93% | $P < 0.001$ | $P < 0.001$ | $P < 0.001$ | $P < 0.001$ | $P < 0.001$ | - |
| Genus | *Blautia* | 3.43% | 3.16% | 4.93% | 2.43% | - | - | - | $P < 0.001$ | $P < 0.001$ | $P < 0.001$ |
| | *Faecalibacterium* | 6.71% | 5.13% | 9.38% | 8.27% | - | - | - | $P < 0.001$ | $P < 0.01$ | $P < 0.001$ |
| | *Prevotella* | 10.27% | 7.67% | 6.02% | 21.12% | - | - | - | - | - | $P < 0.05$ |
| | *Roseburia* | 4.88% | 6.07% | 8.00% | 5.06% | - | - | - | $P < 0.001$ | $P < 0.05$ | $P < 0.001$ |
| | *Subdoligranulum* | 4.57% | 5.90% | 3.56% | 3.32% | - | - | - | $P < 0.05$ | $P < 0.05$ | $P < 0.05$ |
| | *Anaerostipes* | 0.69% | 1.35% | 1.50% | 0.47% | - | - | - | $P < 0.001$ | $P < 0.01$ | $P < 0.001$ |
| | *Escherichia_Shigella* | 0.68% | 2.11% | 0.75% | 0.42% | $P < 0.05$ | - | - | $P < 0.05$ | - | - |
| | *Fusicatenibacter* | 1.34% | 1.50% | 1.68% | 2.05% | - | - | - | $P < 0.001$ | - | $P < 0.001$ |
| | *Haemophilus* | 0.17% | 0.23% | 1.16% | 0.33% | - | - | - | $P < 0.001$ | $P < 0.01$ | $P < 0.001$ |
| | *Lachnospira* | 2.85% | 2.87% | 2.39% | 2.21% | - | - | - | $P < 0.05$ | - | $P < 0.05$ |
| | *Lachnospiraceae_NK4A136_group* | 0.68% | 1.39% | 3.65% | 0.48% | - | - | - | $P < 0.001$ | $P < 0.001$ | $P < 0.001$ |
| | *Megamonas* | 0.54% | 0.95% | 2.77% | 1.01% | - | - | - | $P < 0.01$ | - | $P < 0.01$ |
| | *Phascolarctobacterium* | 1.29% | 2.45% | 2.08% | 1.59% | - | - | - | $P < 0.01$ | - | $P < 0.05$ |
| | *Ruminococcus* | 1.68% | 1.12% | 1.44% | 1.37% | - | - | - | $P < 0.05$ | - | $P < 0.01$ |

**Table 2.** Comparative Analysis of Clinical and Anthropometric Parameters Among Normal, Precocious, Obese, and Obesity-PP Comorbid Groups.

| Taxa | Relative abundance |  |  |  | P value |  |  |  |  |  |
| --- | --- | --- | --- | --- | --- | --- | --- | --- | --- | --- |
|  | Normal<br>(N=18) | Precocious<br>(N=25) | Obes<br>e(N=18) | Obesity-PP<br>comorbid<br>(N=7) | Normal vs<br>Precocious | Normal vs Obesity-PP<br>comorbid | Precocious vs Obesity-PP comorbid | Normal vs<br>Obese | Obesity-PP comorbid<br>vs Obese | Precocious<br>vs Obese |
| Age (years) | 7.61 ± 1.07 | 7.38 ± 0.99 | 6.68 ± 1.05 | 7.41 ± 0.57 | 0.447 | 0.561 | 0.93 | 0.007 | 0.07 | 0.022 |
| Height (cm) | 127.72 ± 6.71 | 125.33 ± 7.76 | 123.74 ± 6.52 | 134.11 ± 1.72 | 0.258 | 0.007 | 0.005 | 0.057 | <0.001 | 0.439 |
| Weight (kg) | 25.66 ± 4.07 | 24.58 ± 4.52 | 29.66 ± 4.92 | 35.81 ± 5.92 | 0.442 | <0.001 | <0.001 | 0.007 | 0.011 | <0.001 |
| BMI | 15.67 ± 1.43 | 15.58 ± 1.29 | 19.31 ± 2.07 | 19.89 ± 2.96 | 0.794 | <0.001 | <0.001 | <0.001 | 0.594 | <0.001 |
| BA (Bone Age) | - | 8.18 ± 1.36 | 7.79 ± 1.29 | 9.46 ± 0.56 | - | - | 0.016 | - | 0.001 | 0.314 |
| BA Advance | - | 0.81 ± 1.14 | 1.44 ± 1.07 | 2.06 ± 0.78 | - | - | 0.005 | - | 0.134 | 0.046 |
| Uterine Volume (cm<sup>3</sup>) | - | 1.38 ± 0.78 | - | 1.15 ± 0.61 | - | - | 0.442 | - | - | - |
| Left Ovarian Volume (cm<sup>3</sup>) | - | 1.58 ± 0.86 | - | 1.84 ± 1.18 | - | - | 0.522 | - | - | - |
| Right Ovarian Volume (cm<sup>3</sup>) | - | 1.63 ± 0.94 | - | 1.81 ± 0.82 | - | - | 0.608 | - | - | - |
| Number of follicles >4mm | - | 6.29 ± 3.76 | - | 5.57 ± 3.41 | - | - | 0.623 | - | - | - |

At the genus level, there were 24, 18, 16, 105, 78, and 104 genera in the in normal and precocious groups, normal and obesity-PP comorbid groups, precocious and obesity-PP comorbid groups, normal and obese groups, obesity-PP comorbid and obese groups, precocious and obese groups.

In species exhibiting higher relative abundance (relative abundance over 1%), the abundance of *Blautia*, *Faecalibacterium*, *Roseburia*, *Anaerostipes*, *Haemophilus*, and *Lachnospiraceae_NK4A136_group* in the obese group significantly higher than that in other three groups (all *P* < 0.05). However, the abundance of *Lachnospira*, *Subdoligranulum* in the precocious group and normal group were significantly higher than that in the obese group, and the abundance of *Prevotella* in the precocious group was also significantly higher than that in the obese group (all *P* < 0.05). Notably, the abundance of *Escherichia_Shigella* in the obese group and precocious group were significantly higher than that in the normal group (all *P* < 0.05). The above results collectively suggested that the microorganisms in the obese group were significantly different from those in the normal group and the precocious group, and that obesity and precocious puberty would significantly increase the level of the harmful bacteria (such as *Escherichia_Shigella*) in the gut.

**It should be specifically noted that the sample size of this study was limited, particularly for the obesity-PP comorbid group (n=7), and as the data did not meet the assumption of normal distribution, the Kruskal-Wallis test was selected to determine significant differences.** While this limits the statistical power for some comparisons, the effect sizes observed for key taxa (e.g., the dramatic increase in Verrucomicrobiota) were substantial and highly significant, suggesting robust biological effects that warrant further investigation in larger cohorts. The above results collectively suggested that the microorganisms in the obese group were significantly different from those in the normal group and the precocious group, and that obesity and precocious puberty would significantly increase the level of the harmful bacteria (such as *Escherichia_Shigella*) in the gut.

## 4. Discussion

This study uncovers specific alterations in the gut microbiota of girls with obesity and precocious puberty (PP), and identifies distinct dysbiotic patterns associated with the comorbidity of these two conditions, highlighting the complex microbiota-host interactions in metabolic and hormonal regulation. Elucidating the underlying mechanisms through which gut microbiota contributes to the pathogenesis of both obesity and PP is essential for translating these findings into targeted microbial interventions. Particularly significant is the discovery of a bidirectional feedback loop involving obesity, estrogen, and Verrucomicrobiota, which provides a novel microbiological perspective for understanding the mechanisms underlying obesity-PP comorbidity.

Importantly, we found the profound alteration of the gut microbiota in obese girls compared to normal girls. While microbial diversity and overall community structure remained similar across all groups, obese girls exhibited significantly increased microbial richness. This suggests that alterations in the gut microbiota, characterized primarily by an expanded repertoire of bacterial species rather than by fundamental changes in community evenness or overall phylogenetic profile, could drive the development of obesity. Notably, this increased richness coincided with a characteristic alteration in the gut microbiota composition at the phylum level, specifically an elevated *Firmicutes*/*Bacteroidota* (F/B) ratio in the obese group compared to both normal and PP groups. This finding aligns with numerous studies linking a higher F/B ratio to obesity (Bervoets et al., 2013; J. Wang et al., 2024). The underlying mechanism may involve the more efficient energy extraction from food by *Firmicutes* compared to *Bacteroidetes*, resulting in enhanced calorie absorption and consequent weight gain (Magne et al., 2020). Additionally, we observed an increase in *Proteobacteria* in the obese group compared to the normal group. Elevated Proteobacteria abundance might promote obesity pathogenesis through lipopolysaccharide (LPS)-mediated inflammation (Zhao et al., 2022). At the genus level, the obese group exhibited significantly higher abundances of *Blautia*, *Faecalibacterium*, *Roseburia*, *Anaerostipes*, *Haemophilus*, and *Lachnospiraceae_NK4A136_group*. It should be noted that some genera (e.g., *Faecalibacterium*) are typically considered beneficial butyrate producers (Maioli et al., 2021). Their increased abundance here, alongside clearly pathogenic groups, suggests complex dysbiosis rather than a uniformly detrimental shift. This phenomenon may be attributed to the fact that in the vast majority of cases—particularly within the intestinal environments of healthy or subhealthy populations—*Faecalibacterium* exerts essential beneficial effects primarily through butyrate production. However, its impact on the host can shift toward harm under altered conditions, such as specific host health statuses, localized chemical milieus (e.g., oxygen availability and inflammatory levels), and shifts in the broader microbial community ecology. This indicates that the metabolic function of *Faecalibacterium* is highly context-dependent: whether it acts as a beneficial or detrimental agent depends critically on the surrounding environmental conditions. This context-dependent duality warrants more in-depth investigation to elucidate the precise mechanisms and ecological determinants governing its functional switch.

Unlike the marked dysbiosis observed in obesity, girls with isolated PP exhibited only minor alterations in gut microbiota composition at the phylum level. The most notable divergence from the normal development group was a significantly reduced relative abundance of Verrucomicrobiota. To our knowledge, previous literature has not reported an association between PP and *Verrucomicrobiota*; thus, the underlying mechanism for this difference remains unclear. Nevertheless, Akkermansia, the primary genus representing *Verrucomicrobiota* in the human gut (Ioannou et al., 2025), provides mechanistic clues. *Akkermansia* is renowned for mucin degradation and its crucial role in maintaining intestinal mucus layer thickness and integrity. It is generally considered a beneficial bacterium associated with enhanced gut barrier function, anti-inflammatory effects, and metabolic health (Cani et al., 2022; Ioannou et al., 2025; Rodrigues et al., 2022). Interestingly, Gu et al. found that epigallocatechin gallate might reverse obesity-induced PP by increasing the abundance of *Akkermansia* (Gu et al., 2024). More epidemiological and functional studies are warranted to investigate the role of *Verrucomicrobiota* and *Akkermansia* in the development of PP. Furthermore, our results indicated that elevated *Escherichia_Shigella* abundance was associated with PP. As Gram-negative organisms, *Escherichia-Shigella* possess LPS in their cell walls. This endotoxin can disrupt blood-brain barrier permeability and promote neuroinflammation (Baske et al., 2024). Research has established associations between *Escherichia_Shigella* and conditions including anxiety disorders (Chen et al., 2019), osteoporosis (Liang et al., 2023), diabetes (Kwan et al., 2022), and obesity (Burakova et al., 2022). However, few studies have investigated the association between gut microbiota and PP, highlighting the need for future research to confirm our findings.

This study reveals that the gut microbiota in the comorbid state of obesity and PP constitutes a distinct ecotype characterized by compensatory adaptations, rather than a simple amalgamation of features from either condition alone. Compared to obesity-only controls, the comorbid cohort exhibited a significant reduction in *Firmicutes* abundance—a notable finding given this phylum’s typical enrichment in obesity. Notably, the comorbidity group exhibited a striking elevation in *Verrucomicrobiota* abundance—approximately 10-fold higher than normal controls (0.46% vs. 5.93%) and nearly 100-fold greater than the obesity-only group (0.06% vs. 5.93%). While increased *Akkermansia* (the primary genus within Verrucomicrobiota phylum) has been documented in gastrointestinal disorders (Y. Zhang et al., 2024) and Parkinson’s disease (Hirayama & Ohno, 2021), *Verrucomicrobiota*’s pronounced upregulation in obesity-PP comorbidity represents a novel observation. The mechanistic basis for this phenomenon remains incompletely understood but may involve obesity-estrogen interactions. Research indicates estrogen promotes the proliferation of *Akkermansia* (Acharya et al., 2019). Given that adipose tissue serves as a significant extragonadal source of estrogen (Hetemaki et al., 2021), estrogenic effects may be amplified in girls with obesity-PP comorbidity, thus manifesting clinically as accelerated expansion of *Verrucomicrobiota* and the dysbacteriosis of microbiota. Given the above evidence, we cautiously propose that *Verrucomicrobiota* may serve as an intervention target for the co-occurrence of obesity and PP in girls. Additionally, the comorbid group displayed reduced microbiota alpha diversity compared to the obesity-only group, potentially attributable to estrogen’s suppressive effect on microbial richness (Acharya et al., 2019). Given the limited research on gut microbiota dynamics in obesity-PP comorbidity, further studies are warranted to validate these findings and elucidate the underlying mechanisms, thereby informing early preventive and therapeutic strategies.

This study has several limitations that should be acknowledged. First, the cross-sectional design precludes causal inferences regarding the relationships between obesity, precocious puberty (PP), and gut microbiota alterations. This limitation is particularly relevant given the well-established bidirectional complexity inherent in gut microbiota–host metabolic interactions. Second, the sample size was limited, especially in the obesity–PP comorbidity subgroup, which may constrain the statistical power and generalizability of our findings. Validation in larger cohorts is therefore required. Additionally, potential confounders such as dietary habits, physical activity, and sleep patterns were not controlled for in the present study. F Given Shenzhen’s status as a migrant city and the complexity/diversity of Chinese dietary habits and cooking methods, controlling for lifestyle factors in human populations would yield poor compliance. Therefore, we plan to utilize animal models in future studies to investigate these relationships under controlled conditions.

Finally, based on the remarkable increase in Verrucomicrobiota abundance observed in the comorbid group in this study, our next step will involve conducting metagenomic and metabolomic analyses of fecal samples from girls with obesity and precocious puberty, as well as blood biochemical tests (including sex hormones). By comparing these data with corresponding indicators from the normal control group, the obesity-only group, and the precocious puberty-only group, we aim to construct a hypothetical model of the bidirectional “gut microbiota-metabolites-hormones” interactions to validate the pathway through which Verrucomicrobiota influences sex hormones. If correlations are observed in the human cohort, animal experiments will be conducted to validate its metabolic functions.

A While 16S rRNA sequencing has provided valuable taxonomic insights, metagenomic sequencing enables a more comprehensive characterization of microbial composition. We plan to conduct metagenomic analyses in subsequent studies and integrate these datasets with metabolomic data.

Finally, the observed elevation in Verrucomicrobia abundance among children with obesity-PP comorbidity suggests a potential bidirectional regulatory mechanism in gut metabolism. We hypothesize that Verrucomicrobia may exert anti-obesity and glucose-lowering effects within a certain range, whereas beyond a specific threshold, it might promote sexual development. Determining the precise critical value for this shift will require further analysis incorporating additional data, such as estrogen (e.g., estradiol) levels from sex hormone assays. Our future work will focus on validating this hypothesis.

## 5. Conclusions

This study demonstrates that the gut microbiota in girls with comorbid obesity and precocious puberty (PP) represents a unique ecotype, distinct from the profiles observed in either condition alone. This specific dysbiotic pattern provides a novel microbiological perspective on the underlying mechanisms of the obesity-PP comorbidity and highlights the potential for targeted microbial interventions.

## Research Funding

This work was supported by grants from the Basic Project of Shenzhen Science and Technology Innovation Commission in 2021,“Discovery of Specific Intestinal Microflora and Intestinal Microecological Intervention for Early Sexual Maturity” (No. JCYJ20210324112804012).

---

**Figure S1.**
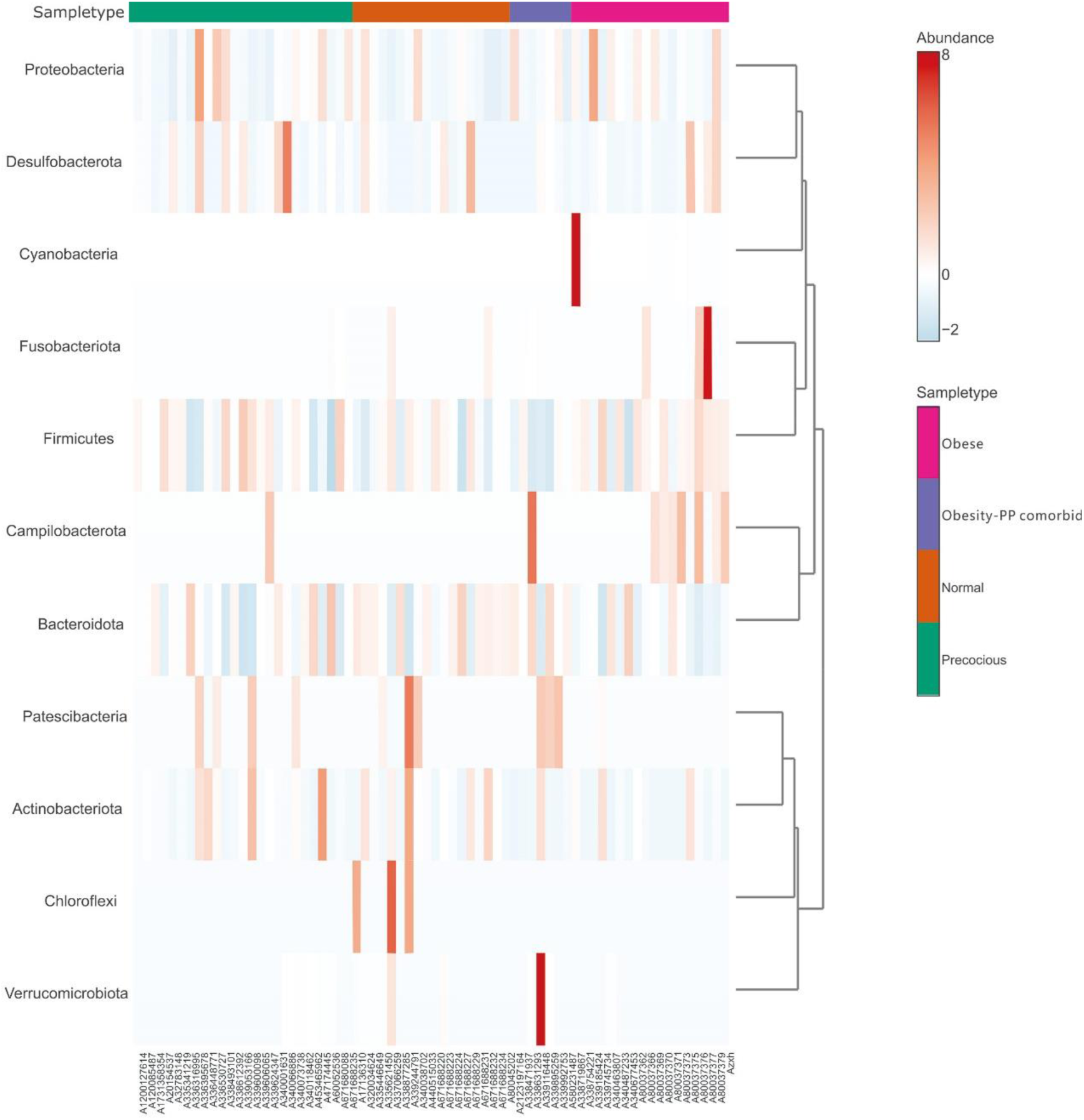
The bacterial composition of gut microbiota at the phylumin different group.

**Table S1.** The sequence number and coverage of 68 samples.

| No. | group | Sequence number | Goods' coverage (%) |
| --- | --- | --- | --- |
| 1 | obese | 32681 | 100 |
| 2 | obese | 34215 | 100 |
| 3 | obese | 36199 | 100 |
| 4 | obese | 37021 | 99.99729883 |
| 5 | obese | 6903 | 100 |
| 6 | obese | 5304 | 100 |
| 7 | obese | 45609 | 100 |
| 8 | obese | 21712 | 100 |
| 9 | obese | 28666 | 100 |
| 10 | obese | 29192 | 100 |
| 11 | obese | 24202 | 100 |
| 12 | obese | 27627 | 100 |
| 13 | obese | 11843 | 100 |
| 14 | obese | 26415 | 100 |
| 15 | obese | 25331 | 100 |
| 16 | obese | 26226 | 100 |
| 17 | obese | 26188 | 100 |
| 18 | obese | 22154 | 100 |
| 19 | obesity-PP comorbid | 7767 | 100 |
| 20 | obesity-PP comorbid | 3841 | 100 |
| 21 | obesity-PP comorbid | 6422 | 100 |
| 22 | obesity-PP comorbid | 7946 | 100 |
| 23 | obesity-PP comorbid | 6307 | 99.9841446 |
| 24 | obesity-PP comorbid | 8148 | 100 |
| 25 | obesity-PP comorbid | 7674 | 100 |
| 26 | normal | 6874 | 100 |
| 27 | normal | 7732 | 100 |
| 28 | normal | 5971 | 99.98325239 |
| 29 | normal | 6644 | 99.98494883 |
| 30 | normal | 6678 | 100 |
| 31 | normal | 5823 | 100 |
| 32 | normal | 9420 | 100 |
| 33 | normal | 8120 | 100 |
| 34 | normal | 5794 | 100 |
| 35 | normal | 4388 | 100 |
| 36 | normal | 3995 | 100 |
| 37 | normal | 7933 | 100 |
| 38 | normal | 6129 | 100 |
| 39 | normal | 5006 | 100 |
| 40 | normal | 7979 | 100 |
| 41 | normal | 3533 | 100 |
| 42 | normal | 5357 | 100 |
| 43 | normal | 4840 | 100 |
| 44 | precocious | 7633 | 100 |
| 45 | precocious | 5783 | 100 |
| 46 | precocious | 4742 | 99.97891185 |
| 47 | precocious | 5222 | 100 |
| 48 | precocious | 6054 | 100 |
| 49 | precocious | 7287 | 100 |
| 50 | precocious | 4139 | 100 |
| 51 | precocious | 10578 | 100 |
| 52 | precocious | 5375 | 100 |
| 53 | precocious | 7900 | 100 |
| 54 | precocious | 7024 | 100 |
| 55 | precocious | 5092 | 100 |
| 56 | precocious | 5642 | 100 |
| 57 | precocious | 7151 | 100 |
| 58 | precocious | 7506 | 100 |
| 59 | precocious | 7286 | 100 |
| 60 | precocious | 8998 | 100 |
| 61 | precocious | 5656 | 100 |
| 62 | precocious | 8684 | 100 |
| 63 | precocious | 6439 | 100 |
| 64 | precocious | 8203 | 100 |
| 65 | precocious | 6259 | 100 |
| 66 | precocious | 6121 | 100 |
| 67 | precocious | 3996 | 100 |
| 68 | precocious | 5454 | 100 |

## Notes

https://meta.bgi.com/microbe/login

### Competing Interest Statement

The authors have declared no competing interest.

